# Bacterial transformation buffers environmental fluctuations through the reversible integration of mobile genetic elements

**DOI:** 10.1101/557462

**Authors:** Gabriel Carvalho, David Fouchet, Gonché Danesh, Anne-Sophie Godeux, Maria-Halima Laaberki, Dominique Pontier, Xavier Charpentier, Samuel Venner

## Abstract

Horizontal gene transfer (HGT) is known to promote the spread of genes in bacterial communities, which is of primary importance to human health when these genes provide resistance to antibiotics. Among the main HGT mechanisms, natural transformation stands out as being widespread and encoded by the bacterial core genome. From an evolutionary perspective, transformation is often viewed as a mean to generate genetic diversity and mixing within bacterial populations. However, another recent paradigm proposes that its main evolutionary function would be to cure bacterial genomes from their parasitic mobile genetic elements (MGEs). Here, we propose to combine these two seemingly opposing points of view because MGEs, although costly for bacterial cells, can carry functions that are point-in-time beneficial to bacteria under stressful conditions (e.g. antibiotic resistance genes under antibiotic exposure). Using computational modeling, we show that, in stochastic environments (unpredictable stress exposure), an intermediate transformation rate maximizes bacterial fitness by allowing the reversible integration of MGEs carrying resistance genes but costly for the replication of host cells. By ensuring such reversible genetic diversification (acquisition then removal of MGEs), transformation would be a key mechanism for stabilizing the bacterial genome in the long term, which would explain its striking conservation.

## Introduction

Horizontal gene transfer (HGT), i.e. the passage of heritable genetic material between organisms by means other than reproduction, is commonly observed in bacteria ^1–3^. By promoting the spread of genes of antibiotic or heavy metal resistance and virulence factors, HGTs weight in an important threat on human health ^4^. Among all known mechanisms by which HGTs occur, one can distinguish those resulting from the infectious and propagative behavior of mobile genetic elements (conjugation, transduction) and those that are exclusively controlled by the bacterial cells ^5–7^. By far, the most widespread of those is natural transformation, i.e. the import of free extracellular DNA (eDNA) and its integration into the bacterial genome by homologous recombination ^8^. The DNA import system is expressed under the state of competence which is triggered by signals that are often elusive and difficult to reproduce under laboratory conditions ^9,10^. Despite this difficulty, transformation has been experimentally demonstrated in over 80 bacterial species distributed throughout the tree of life, indicative of an ancestral origin ^8,9,11^. The list of transformable species keeps growing, now including species that had long been considered incapable of transformation ^12^. In addition, from the recent database AnnoTree grouping nearly 24,000 bacterial genomes ^13^, we uncover that transformation-specific genes required for the uptake of eDNA (*comEC, dprA*) are extremely conserved (Fig. 1). This suggests that most bacterial species may undergo transformation in their natural habitat. In spite of this ubiquity and that this mechanism has been documented for a long time ^14^, the evolutionary causes of transformation are complex to disentangle and are still debated.

**Figure 1:**
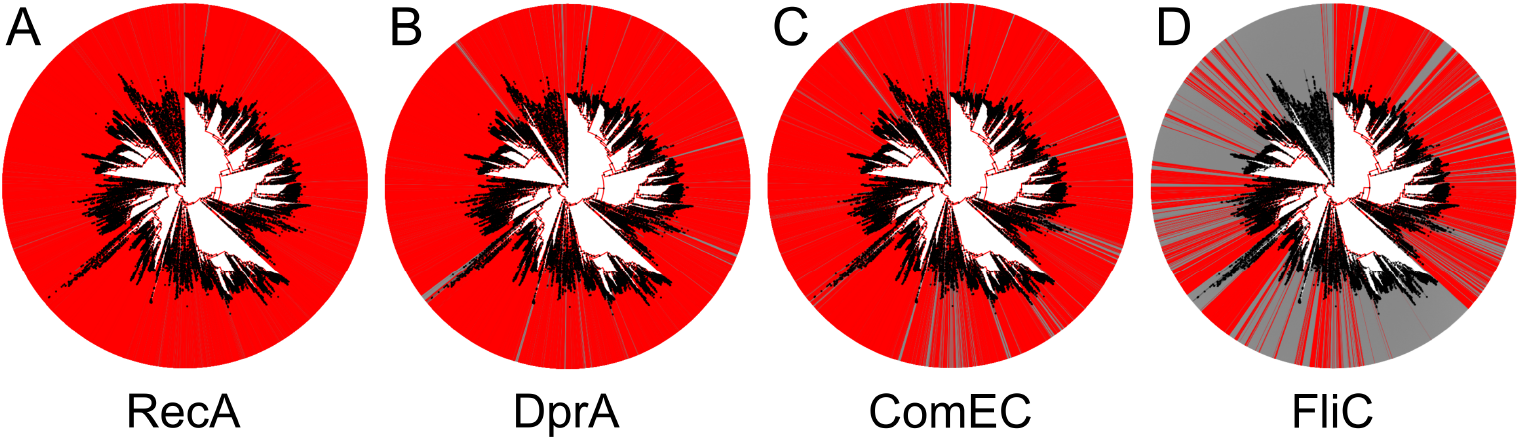
Proportion of competence genes in the AnnoTree database. Graphs represent the phylogenic tree of the 23,936 bacterial genomes. Red lines indicate the gene has been found in the genome. (A) *recA* (KEGG identifier K03553), 23059 genome hits; *recA* is a gene involved in homologous recombination, a core function essential to chromosome maintenance, and is well known for its ubiquity in all bacterial genomes. (B) and (C) are genes required for transformation, specifically expressed during competence, and are therefore indicators of the ability to perform natural transformation: *comEC* (KEGG identifier K02238), 20171 genome hits, involved in the uptake of extracellular DNA and (C) *drpA* (KEGG identifier K04096), 21523 genome hits, involved in protection of the incoming DNA and its integration in the recipient genome. DprA acts by directly and specifically interacting with ssDNA imported through the ComEC channel and then brings to RecA for recombination. Involvement of DprA in processing of ssDNA other than transforming DNA have so far been excluded (generic recombination). (D) In contrast, the distribution of fliC, (KEGG identifier K02406), 9299 genome hits, encoding the subunit of the bacterial flagellum, is consistent with an accessory function.

The commonly accepted evolutionary benefits of transformation is the genetic diversification and mixing within bacterial populations. Computational modeling approaches have shown that gene acquisition and genetic mixing from transformation can provide a selective advantage by allowing bacteria to combine favorable mutations (Fisher-Muller effect) ^15,16^ and to efficiently exploit new or fluctuating environments ^17^. Under fluctuating selection of different alleles, transformable bacteria may also benefit from the acquisition of old alleles present in their environment to restore a fitter phenotype ^18^. In complement to these theoretical investigations, an increasing number of recent studies, in part fueled by the exponential growth of genome sequencing, shows that through transformation bacteria frequently acquire new functions carried by transposons, integrons and genomic islands ^19–22^. Transformation enables acquisition of antibiotic resistance by *Campylobacter jejuni* and capsule switching by *Streptococcus pneumoniae* leading to vaccine escape ^23–25^. *Bacilus subtilis* presents a large accessory genome, a diversity seemingly generated by transformation, allowing this species to colonize various ecological niches, from soils and plants to guts ^21^. Overall, the extracellular DNA obtained from transformation may provide habitat specific genes and favors adaptation to new environments ^26^. Yet, genome-based evidence of the benefit of transformation are inherently biased as they tend to highlight HGT events that result in the acquisition of genes providing clear selective advantage (e.g. antibiotic resistance).

In opposition to the genetic diversification and mixing paradigm, Croucher *et al*. (2016)^27^ recently proposed the radically distinct hypothesis that the main evolutionary function of transformation is to cure bacterial genomes of integrated genetic parasites. Based on the observation that bacterial genomes are inevitably parasitized by MGEs ^28,29^, they argue that transformation favors the insertion of short DNA sequences from kin cells, thus tending to remove infesting DNA rather than to acquire additional foreign DNA ^27,30^. Following this “chromosomal curing” hypothesis ^27^, transformation would be a defense mechanism against genetic parasites rather than a mechanism to generate genetic diversity and mixing. In favor of this proposal, several observations point out that cells are much more likely to undergo recombination with closely related neighbors than with distant lineages carrying foreign DNA ^30,31^. Bacteria usually grow close to their siblings and the emergent physical vicinity favors the exchange of DNA sequences between closely related genotypes. In addition, bacteria may regulate the competence state using quorum sensing, hence ensuring that a critical amount of kin cells are nearby ^32^. They may also select DNA with specific uptake sequences to avoid integrating exogenous DNA ^33^. In some cases, competent bacteria kill their own kin cells, enriching their surrounding with clonal DNA ^34^. Considering all the barriers to the acquisition of foreign DNA, transformation would mainly act as a conservative mechanism instead of a mean of genetic diversification and mixing ^31^.

In this work, we develop a model to combine the apparent two antagonistic points of view on the evolutionary causes of transformation in bacteria. We propose that the combination of both roles, gene acquisition and chromosomal curing, allows transformable cells to reversibly integrate MGEs that carry resistance genes which would be of particular interest in stochastically fluctuating environments. This proposition relies on the fact that (i) bacteria are exposed to a large variety of stochastic stresses ^35^, (ii) genes conferring resistance to these stresses (e.g. antibiotics and heavy metals) are often carried by MGEs such as genomic islands and not in the core genome ^36,37^ and finally (iii) MGEs carrying resistance genes are costly for the replication of their host and are not required in stress-free environments ^38^. Using a computational model introducing competition between bacterial genotypes with different transformation strategies (transformation rate) and stochastic stressful environments, we show that intermediate transformation rates are optimal in fluctuating environments. Natural transformation stabilizes the bacterial genome by allowing both the acquisition and cure of MGEs carrying resistance which would only be unnecessary genetic parasites in stress-free environments.

## Results

### Transformable cells are competitive in stochastic environments

Based on the model summarized in Fig. 2 and detailed in the method section, we simulate competition between several bacterial genotypes differing in their transformation rates. Population dynamics are simulated in four distinct environments: one stress-free constant environment and three environments with stochastic stress exposure, differentiated by stress frequency (Supplementary Figure 1). In each tested environment, bacterial genotypes share the same eDNA pool resulting from bacterial dead cells and initially composed of wild type (WT) alleles. Small amount of MGEs carrying a stress resistance gene are introduced to this pool. The transformable genotypes also compete with two control genotypes: a non-transformable genotype susceptible to the stress (NTS) and a non-transformable resistant genotype (NTR) carrying a resistance gene that has the same cost in terms of cell replication as the MGE (see methods).

**Figure 2:**
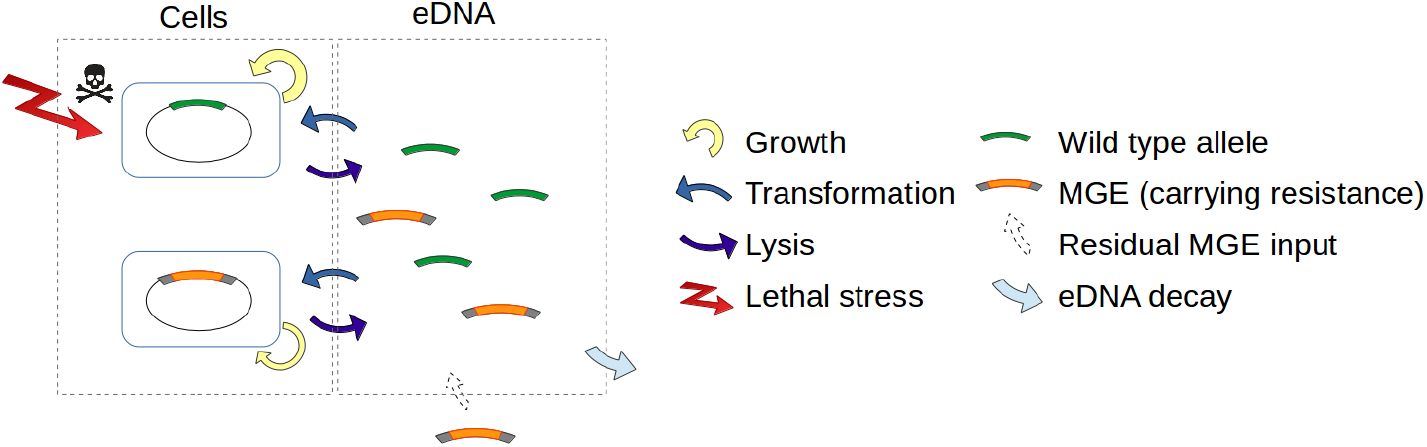
Schematic representation of the computational model. Bacterial cell growth follows a logistic growth model. The bacterial cells have an insertion site in their chromosome at which two types of alleles from the eDNA compartment can be integrated from transformation and replace their current DNA: wild type (WT) allele and MGE. The integration of a WT allele is costless for cells whereas the integration of MGEs causes a cost in terms of cell replication. Bacterial populations are faced with stochastic stresses of random duration and intensity. In the absence of stress, cells are lysed at a basal rate. Under stress exposure, the lysis rate of WT cells increases but remains unchanged for cells with a MGE carrying resistance. Each lysed cell releases its DNA and fuels the extracellular compartment with eDNA. MGEs are constantly added to the extracellular environment at a marginal rate (MGE input) simulating a residual arrival from neighboring populations. The WT alleles and MGEs are degraded at a constant rate in the extracellular compartment.

In the stress-free environment, all genotypes can grow when they are alone (Supplementary Figure 2). In the competition context, the NTR genotype becomes extinct because carrying a resistance gene reduces growth rate, while all other genotypes have similar demographic performances and do not experience extinction regardless of their transformation rate (Fig. 3A, 3B). This is related to the fact that there is no transformation cost in these simulations (see Supplementary Figure 3, 4 when a cost is introduced). As MGEs carrying resistance are extremely marginal in the extracellular compartment (Fig. 3D), therefore upon transformation, cells mostly integrate WT alleles into their genome and conserve their wild type phenotype (Fig. 4A).

**Figure 3:**
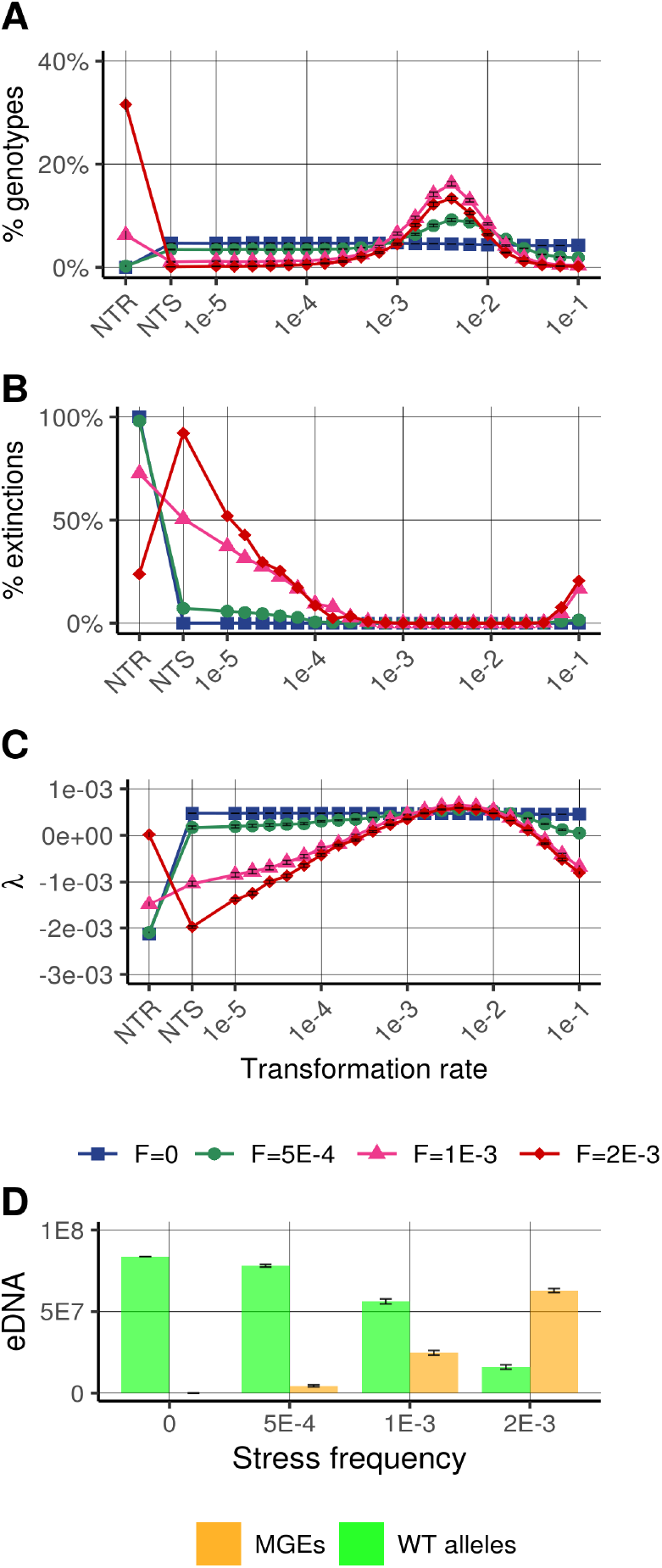
Relative success of genotypes according to their transformation strategies. (A) Proportion of the competing genotypes at *t*=5000: non-transformable resistant (NTR), non-transformable susceptible (NTS) and 21 genotypes with transformation rates ranging from 10^−5^ to 10^−1^TU^−1^. (B) Proportion of extinction of the genotypes at *t*=5000, among the 500 simulations. (C)Stochastic growth rate, as proxy of the fitness of the genotypes in stochastic environments (see methods). (D) Composition of the extracellular compartment at *t*=5000. Represented data is the mean and standard error calculated form 500 simulations. Population dynamics are simulated in four distinct environments: one stress-free constant environment (F=0) and three environments with stochastic stress exposure, differentiated by stress frequency of F=5*10^−4^, F=10^−3^ and F=2*10^−3^ (Supplementary Figure 1).

**Figure 4:**
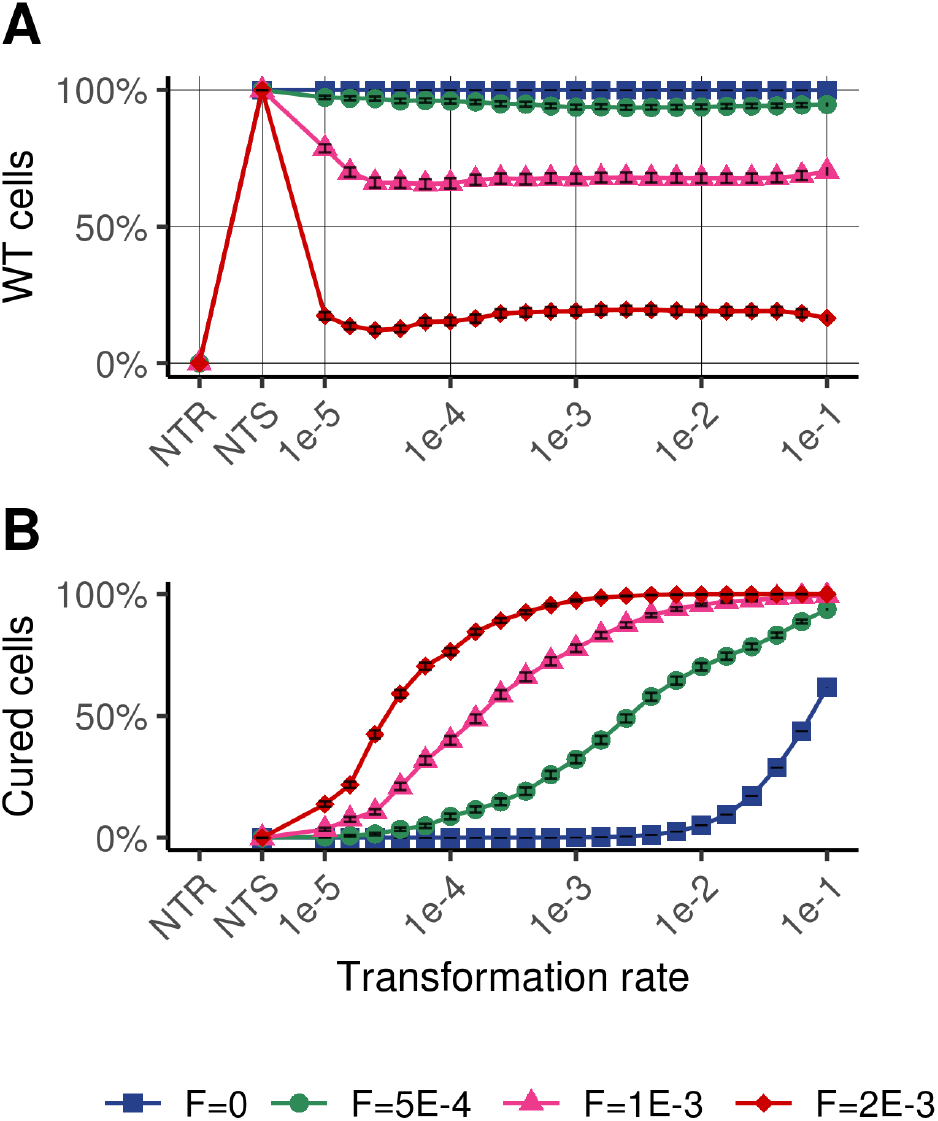
Maintenance of wild type alleles in transformable genotypes. (A) Percentage of wild type (WT) cells at the end of the simulations for each genotype. Extinct genotypes are not accounted for the calculus. (B) Percentage of WT cells from cure, i.e. WT cells with past MGE integration.

When bacterial cells are exposed to stress, even at a low frequency, the most efficient strategies that emerge are those with an intermediate transformation rate (10^−3^<*T_max,i_*<10^−2^) which have both a high abundance, no extinction and a higher stochastic growth rate (Fig. 3A, 3B, 3C). The NTS genotype, which cannot acquire resistance genes, is disadvantaged and often goes extinct, whereas it could grow if it are alone, i.e. without competition with other genotypes (Supplementary Figure 2). The performance of genotypes with a very low transformation rate (*T_max,i_*<10^−4^) is very similar to that of the NTS genotype even if the extinction probabilities of these genotypes are lower (Fig. 3A, 3B). Interestingly, and even without introducing a direct cost for transformation (in terms of replication or cell mortality), cells that transform at a very high rate (*T_max,i_*>10^−2^) are also counter-selected (Fig. 3A, 3B). These strategies induce very frequent changes in phenotype, including the detrimental transitions WT → resistant (MGE infected) between stresses and resistant (MGE infected) →WT during stresses (Supplementary Figure 5).

Although the NTR genotype always perform well alone (Supplementary Figure 2), it is outcompeted in all environments except in few simulations with high stress frequency (2x10^−3^) (Fig. 3A). This result shows that continuously carrying the resistance gene may be beneficial if the environmental stress is relatively frequently encountered. However, despite that in the most stressful environment the NTR genotype has the highest mean total cells at the end of the simulations it actually suffers ~25% extinction (percentage that increases with simulation time, Supplementary Figure 6), whereas the predominant transformable genotypes persist in all simulations (Fig. 3B), and have a higher stochastic growth rate than NTR genotypes (Fig. 3C). The observation that the intermediate transformable genotypes outcompete the NTR genotype suggests that the cure of the genome gives a fitness advantage in stochastic environments by removing MGEs, costly for replication, during the periods without stress.

The results obtained are robust, even if we consider the inherent risks associated with transformation: the possibility of acquiring toxic and highly detrimental genes and the recombination intermediates that can jeopardize chromosome integrity ^39,40^. To account for such a transformation cost, we implemented a probability of cell lysis during transformation events. By including this cost (up to 10% risk of death during transformation), genotypes with high transformation rates are greatly impaired in the stress-free environment (Supplementary Figure 3). However, in stressful environments, strategies with intermediate transformation rates (10^−3^<*T_max,i_*<10^−2^) remain optimal, even if the optimum shifts towards the lower transformation rates (Supplementary Figure 4). Similarly, when the cost of carrying resistance is modified, the cost reduction favors the NTR genotype, while the cost increase still favors intermediate transformation rates (Supplementary Figure 7). Moreover, the variation in the input of MGEs or in the rate of degradation of eDNA does not qualitatively change the results (Supplementary Figure 8, 9), nor the specific accentuated degradation of MGEs (up to 10 times the one of WT alleles) (Supplementary Figure 10). Finally, the optimal intermediate transformable genotypes remain stable when we modify the mean intensity or mean duration of the stresses, although the NTR genotype suffers great variations and constantly higher extinctions (Supplementary Figures 11, 12). Overall, results point out that intermediate transformation rates are extremely efficient strategies for buffering environmental stochasticity in many ecological contexts.

### The competitiveness of transformable cells relies on the reversible integration of MGEs

To examine the importance of chromosomal curing in the success of genotypes with intermediate transformation rates (Fig. 3ABC, transformation rates 10^−3^<*T_max,i_*<10^−2^), we determined the phenotypic composition (proportion of WT cells and cells infected by MGEs) for the different genotypes (Fig. 4A) as well as the proportion of WT cells from parent cells that previously had a MGE in their genome (WT cells from cure, Fig. 4B). The proportion of WT cells is similar in all transformable genotypes despite genotypes being represented in various proportions (Fig. 3A). The proportion of WT cells, however, decreases as the stress frequency increases. By determining the origin of WT cells in stochastic stressful environments, we found that most of them originate from genome cure for the predominant genotypes (Fig. 4B, transformation rates between 10^−3^ and 10^−2^). The proportion of WT cells from cure increases with the frequency of stress exposure (Fig. 4B) and with the simulation time (Supplementary Figure 6).

These results show that transformation, performed at an intermediate rate, is a powerful mechanism for regenerating the WT genotype in stochastic environments. From the analysis of transformation events performed by the dominant genotype (*T_max,i_*=10^−2.4^), we find that the switch of phenotypes (WT → resistant or resistant → WT) occurs mainly when the genotype faces intermediate numbers of stress during simulations (~4 to 7 stresses) (Fig. 5A, 5B), which also corresponds to a greater diversity in the composition of the extracellular compartment (Fig. 5C). With extreme number of stresses, the environment is either very little disturbed or, conversely, frequently disturbed and the great majority of transformation events are neutral since they replace the DNA of the cells with another identical DNA (Fig. 5).

**Figure 5:**
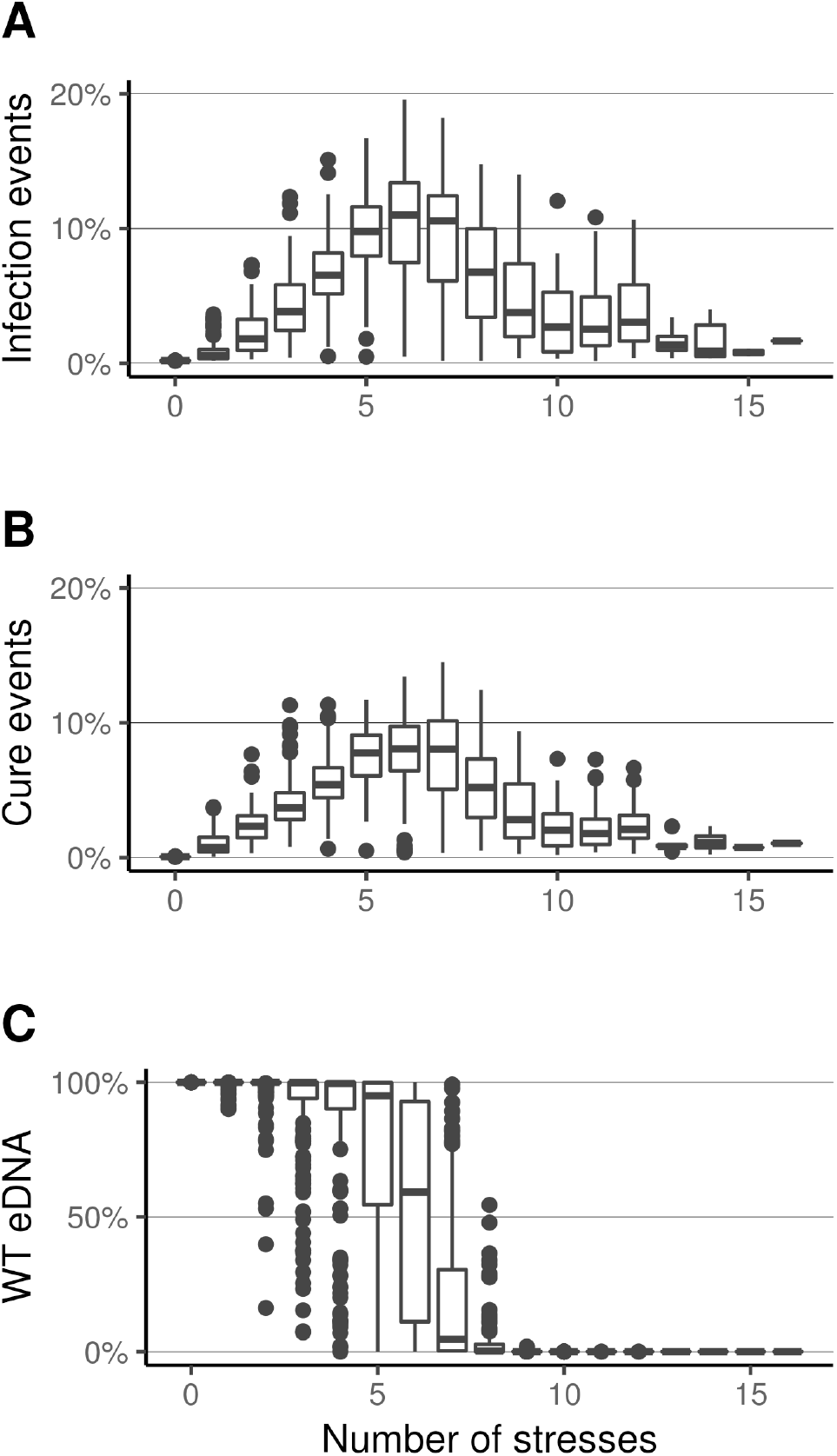
Types of transformation events (infection or cure) for the predominant genotype (with transformation rate 10^−2.4^). (A) Box plot of the percentage of cure events (MGE infected cell→WT cell) among all transformation events depending on the number of stresses. (B) Box plot of the percentage events of infection by MGE (WT cell → MGE infected cell) among all transformation events depending on the number of stresses. Results groups simulations with all stress frequencies. (C) Composition of the extracellular compartment at *t*=5000 (Box plot of the percentage of WT alleles).

## Discussion

While transformation is often considered as a mechanism sustaining bacterial genetic diversification and mixing, recent investigations suggest that its primary function would be to cure bacterial genomes of their parasitic genetic elements ^27,30^. We show that these two points of view can be unified in a common framework. In our simulations, genotypes with an intermediate transformation rate, when competing with other genotypes, have a large selective advantage in fluctuating environment (stochastic stress exposure) by maximizing the probability of genotype persistence and their stochastic growth rate (Fig. 3B, 3C). In our simulations, these intermediate transformation rates favor the random acquisition of extracellular MGEs carrying resistance genes, which allows genotypes to continue to grow during periods of environmental stress. Furthermore, these same transformation strategies allow the MGE removal (genome cure) after each stress episode and thus the reconstitution of the initial genome, which is beneficial because maintaining MGEs is costly in terms of replication for host cells ^36^. Overall, our results suggest that intermediate transformation rates, by generating reversible integration of MGEs, stabilize over the long term the genotypes and genomes of bacteria that evolve in a stochastic environment.

Bacterial populations often face variable environments and are exposed to a wide variety of unpredictable stresses (e.g. heavy metals, antibiotics). The most widely proposed adaptation mechanisms to deal with them correspond to the diversified bet hedging - stochastic switching between phenotypic states ^41–44^ - corresponding to a risk-spreading strategy that facilitates genotype invasion and persistence in the face of unpredictable fluctuating environmental conditions ^45^. The most common example corresponds to the production of both reproductive individuals (replicative cells in bacteria) and individuals that remain in a dormant state for a more or less prolonged period of time (or “persister” cells in bacteria) ^46,47^. Similarly, transformation can be seen as a risk-spreading strategy by randomly producing new phenotypes from MGEs integration, then reconstituting the initial genotype from MGE removal. Interestingly, this strategy would enable to cope with a wide variety of stresses (by successively integrating and removing different MGEs) while maintaining active (replicative) cells during and between periods of stress. In this sense, transformation could be one of the most efficient risk spreading strategies in stochastic environment, which could explain its ubiquity in the bacterial phylum (Fig. 1).

When in competition, genotypes with transformation rates too low or too high should be counter-selected because they generate too little or too much phenotypic change, often leading to the formation of phenotypes that are not adequate for environmental conditions (cells with MGE in stress-free period or WT cells during exposure to stress, Supplementary Figure 5). Mutants displaying phenotypes of elevated transformability are often isolated in transformable species under laboratory conditions^48–52^. Such hypertransformable phenotype is due to mutations causing upregulation of the transformation system or alteration of component of the machinery. Interestingly, these hypertransformable phenotypes are not common in natural isolates, suggesting that they can arise but are quickly counter-selected. Our prediction is consistent with the fact that in many transformable species, only a fraction of cells of the same genotype is in the state of competence and are likely to transform at some time, even if they are placed under favorable controlled laboratory conditions ^8,53–55^. This diversification of competence states, which would correspond to the intermediate transformation rate in our model, should greatly contribute to the spreading of risks in a stochastic environment.

According to our proposal, the extracellular compartment would constitute a reservoir of MGEs, providing bacteria with a “communal gene pool” ^56^, which should be highly fluctuating in its composition. In our model, we introduce MGEs at an extremely low rate, simulating the residual intake of MGEs from other nearby bacterial populations. The proportion of these MGEs remains extremely low when stress is rare or absent (Fig. 3D) while they can become extremely abundant in the extra-cellular compartment when stress exposure become frequent (Fig. 3D). Our work then encourages to empirically analyze the dynamics of the extracellular compartment whose composition should depend on the regime of environmental fluctuations (intensity, duration, frequency of exposure to stress), and the degree of persistence of MGEs or wild alleles according to their characteristic (e.g. their size). With the advent of methods to study bacteria-MGE interactions in complex microbiota ^57,58^, contrasting antibiotic treatments could be an ideal experimental design to explore the dynamics of this extracellular reservoir and its consequences on the spread of resistance genes. For example, permanent antibiotic treatment could enrich the extracellular compartment with MGE to such an extent that bacteria can no longer cure their genome through transformation. This point of view shed new light on our understanding of (and fight against) the spread of antibiotic resistance in hospitals and potentially pave the way to new strategies for fighting antimicrobial resistance.

The most common hypothesis that transformation is exclusively a means of generating genetic diversification faces various obstacles (see introduction). However, there are undoubtedly empirical facts in favor of the perennial acquisition of new genes from different species by natural transformation ^59,60^. We propose that such integration should be the result of rare and “accidental” transformation events leading to the formation of new bacterial strains potentially in competition with the parent strain. Therefore, one of the main evolutionary causes of transformation should be to generate reversible integration of MGE to buffer environmental stochasticity, while the perennial acquisition of new genes should be a by-product of transformation only exceptionally occurring. These accidental transfers would however play a key role in the diversification of bacterial lines and would be of the same order as horizontal transfers observed in eukaryotes, which are rare but have a profound impact on their evolution ^61,62^.

On the opposite to the genetic diversification hypothesis, the “chromosomal curing” hypothesis disregards the acquisition of new genes from transformation and focuses on the removal of MGEs, often view as parasites of bacterial genomes ^27^. This point of view is particularly relevant in the case of MGE phages which are infectious and, according to the current state of knowledge, very rarely carry any resistance genes ^37^. However, the “chromosomal curing” hypothesis alone cannot explain the frequent observation of accumulation of resistance genes in MGEs, such as resistance islands which can be very numerous and diversified ^36,63^. Here, we propose that bacteria, through transformation, actively exploit the MGEs of the extracellular compartment. In return, MGEs must be seen as entities competing for the exploitation of the bacterial resource and co-evolving with bacteria. In this line, an efficient strategy for MGEs could be to interfere with transformation ^20^, which could reduce their risk of being eliminated from bacterial genomes.

In conclusion, transformation, which is an extremely well conserved trait, may be one of the most widespread mechanisms by which bacteria can cope with environmental stochasticity. The reversible integration of certain MGEs would then allow bacteria to stabilize their genome in the long term. Understanding the evolution of bacterial populations and communities in a fluctuating environment will require addressing the co-evolution of bacteria and MGEs by considering a possible alternation of genetic conflicts and cooperation between them. This new perspective could help understand and prevent the spread of antibiotic resistance in bacterial populations and communities.

## Methods

We developed a stochastic computational model that included two types of compartments: bacterial cells and extracellular DNA (eDNA), similarly to previous models ^27^. The overall structure of the model is displayed in Fig. 2. Bacterial cells have an insertion site in their chromosome which can be occupied by two DNA types: wild type (WT) allele and costly MGE conferring stress resistance. In the population, 23 genotypes *i* compete with each other’s among which 21 genotypes *i* differ in their transformation rate. We also introduced two control non-transforming genotypes, one with the WT allele and one with the stress resistance allele. The NTR genotype is initialized with cells carrying the stress resistance allele while the transformable and the NTS genotypes are initialized with WT cells only, i.e. cells carrying the WT allele. Genotypes, according to their transformation strategy, are equitably distributed at the beginning of the simulations, with an initial population size *N*_0_=10^5^ cells. Bacterial growth follows a logistic growth model, with a carrying capacity *K*=10^7^cells (i.e. the maximum number of cells that the habitat can support). The number of replicating cells per genotype *i* and per time step *dt, G_i,t+1_*, is determined using a binomial distribution:

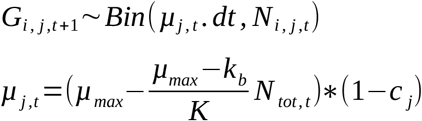

*N_i,j,t_* is the number of cells with the genotype *i* carrying the DNA type *j* at time *t. μ_j,t_* is the replication rate of cells with the DNA type *j* at time *t. μ_max_* is the maximal growth rate. *k_b_* is the basal lysis rate, independent of the presence of stress. *N_tot,t_* is the total number of cells in the population (considering all genotypes) at time *t* and *c_j_* is the cost induced by the DNA type *j* on the replication of cells (*c_WT_*=0 and *C_MGE_*>0). The number of lysed cells per time step *L_i,j,t+1_* is determined using a binomial distribution:

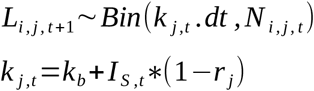

*T_j,t_* is the lysis rate of cells with the DNA type *j* at time *t*. *I_S,t_* is the intensity of the stress at time *t*. Stresses increase the lysis rate of cells carrying a wild type allele whereas the lysis rate of cells carrying a MGE remains at the basal rate *k_b_* (*r_WT_*=0 and *r_MGE_*=1). The number of competent cells undergoing a transformation event during a time step *C_i,t+1_* is determined using a binomial distribution:

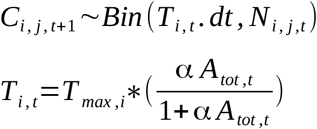

*T_i,t_* is the transformation rate at time *t*. *T_max,i_* is the maximal transformation rate of the genotype *i* and is the only parameter differentiating the transformable genotypes. *α* is the binding rate between cells and eDNA. *A_tot,t_* is the total number of eDNA at time *t*. The probability of a cell to take up a particular type of eDNA is proportional to the DNA composition of the extracellular compartment. Cells undergoing a transformation event change their genotype accordingly to the DNA type integrated. The overall variation of a genotype *i* containing a DNA type *j* (WT or MGE) during a time step is summarized by:

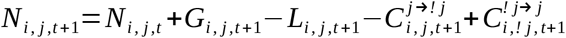

In the extracellular compartment, eDNA is degraded at a constant rate *R*. The number of degraded eDNA molecules per time step *D_j,t+1_* is determined using a binomial distribution:

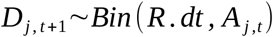

*A_j,t_* is the number of eDNA of type *j*. The extracellular compartment is alimented by eDNA from lysed cells, each lysed cell releasing a DNA molecule corresponding to their DNA. In addition, eDNAs are added at a constant rate to the extracellular compartment. The number of eDNA molecules added per time step *M_j_* is defined by:

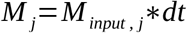

*M_input,j_* is the number of molecules of eDNA of type *j* added per time unit. Only MGEs are added this way in the extracellular compartment (*M_input,WT_*=0). The overall variation of a DNA of type *j* in the extracellular compartment during a time step is summarized by:

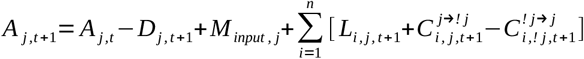

Bacteria are exposed to random stresses affecting the lysis rate of WT cells and occurring with a frequency *F_Stress_*. The probability of a stress to start during a time step is *F_stress_*dt*. When a stress starts, another one cannot begin before the first one ended. The duration *d_S_* and intensity *I_S_* of each stress is set randomly drawn from a normal distribution with the means and standard deviations *d_mean_, d_sd_, I_mean_ and I_sd_*. The summary of the parameters used is presented in Table 1. The stochastic growth rate λ is calculated as the logarithm of the geometric mean:

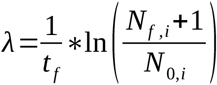

with *t_f_* the simulation time, *N*_*f*,i_ the final number of cells of the genotype *i* and *N_0,i_* the initial number of cells of the genotype *i*.

**Table 1:**
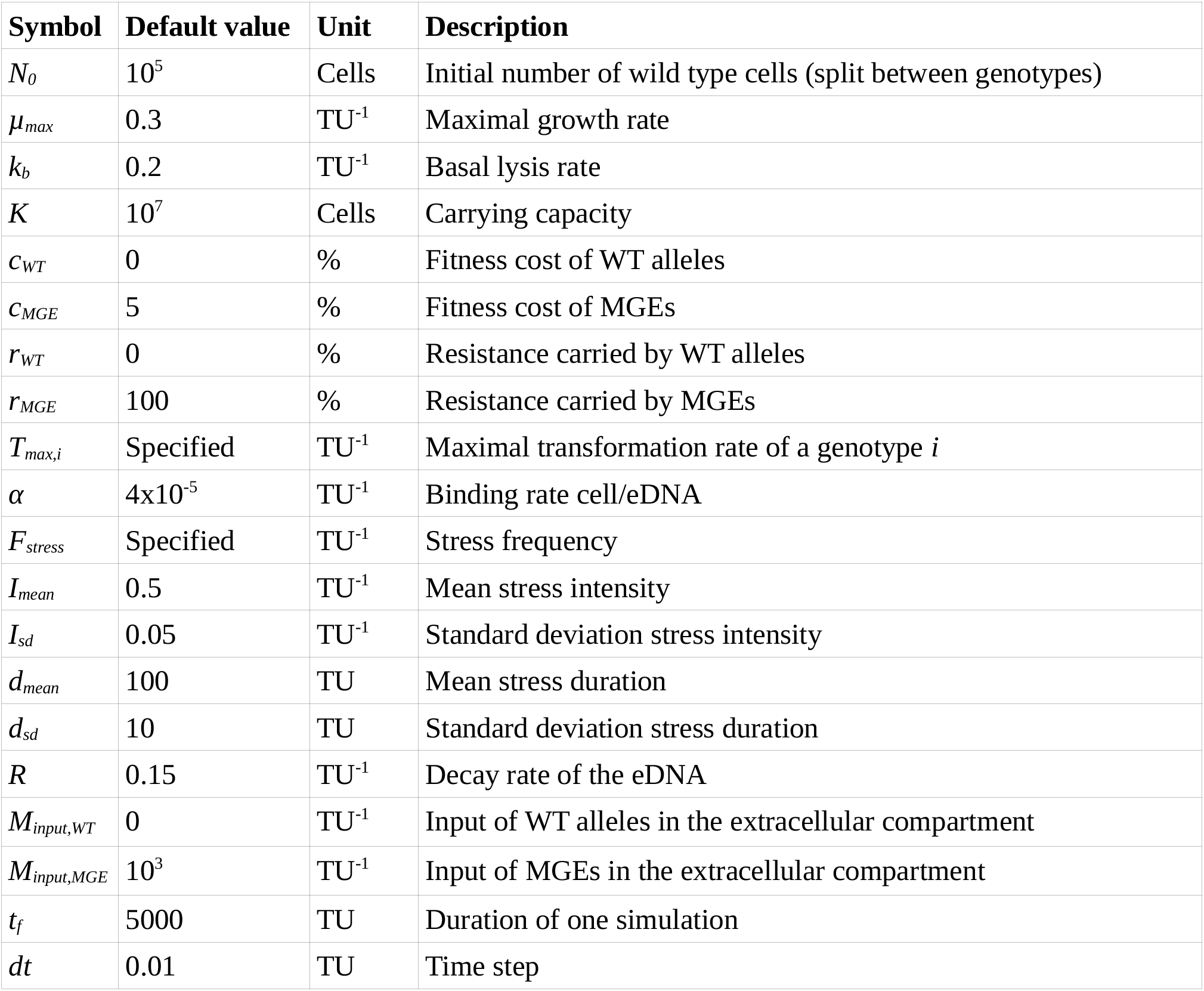
Default parameters used. (TU=Time Unit)

## Code availability

The C++ code is available by requesting the authors.

## Supporting information

Supplemental Table anf Figures

## Acknowledgments

We warmly thank Vincent Miele for his computer programming advice. This work was supported by the LabEx ECOFECT (ANR-11-LABX-0048) of Université de Lyon, the Centre National de la Recherche Scientifique (CNRS) and the Université de Lyon 1. Simulations were performed using the computing cluster CC LBBE/PRABI.

## Author contributions

SV and XC conceived and led the study; SV, XC, DF, GD, GC design the model; GD and GC program the model; GC manages the simulation program; GC, SV, XC wrote the paper; All authors gave fruitful comments during the research process and revised the manuscript.

