## Supplemental Table anf Figures for "Bacterial transformation buffers environmental fluctuations through the reversible integration of mobile genetic elements"

### Supplementary information

Gabriel Carvalho[1], David Fouchet[1], Gonché Danesh[1], Anne-Sophie Godeux[2], Maria-Halima Laaberki[2,3], Dominique Pontier[1], Xavier Charpentier[2, 4,*], Samuel Venner[1, 4, *]

[1]Laboratoire de Biométrie et Biologie Evolutive, CNRS UMR5558, Université Claude Bernard Lyon 1, Villeurbanne, France
[2]CIRI, Centre International de Recherche en Infectiologie, Inserm, U1111, Université Claude Bernard Lyon 1, CNRS, UMR5308, École Normale Supérieure de Lyon, Univ Lyon, 69100, Villeurbanne, France
[3]Université de Lyon, VetAgro Sup, 69280 Marcy l'Etoile, France.

[4] These authors contributed equally to this study

Simulations are run with the standard parameters (main text, Table 1) and the stress frequency $10^{-3}TU^{-1}$ unless specified.

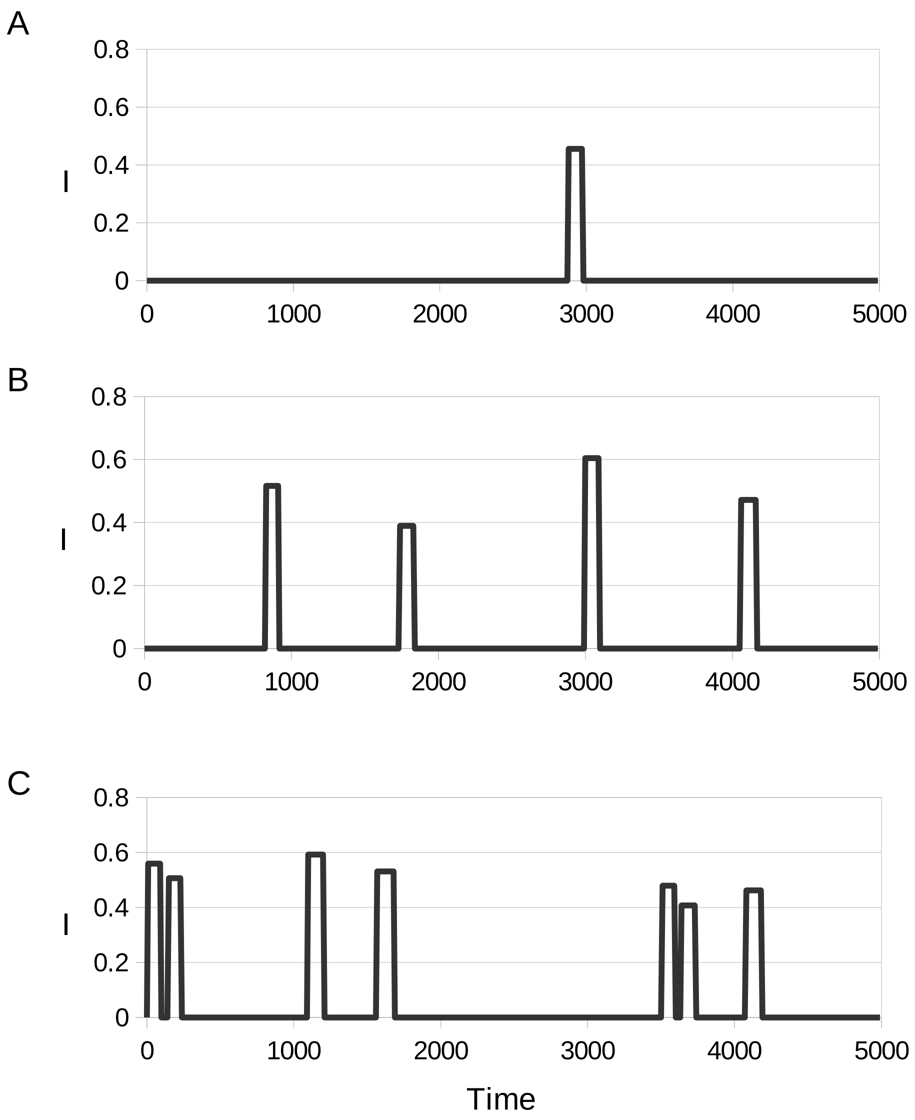


*Sup. Figure 1:* Examples of stress dynamics in the three environments with stochastic stress exposure. I is the intensity of the stresses over time for a stress frequency of $5*10^{-4}$ (A), $10^{-3}$ (B) and $2*10^{-3}$ (C).

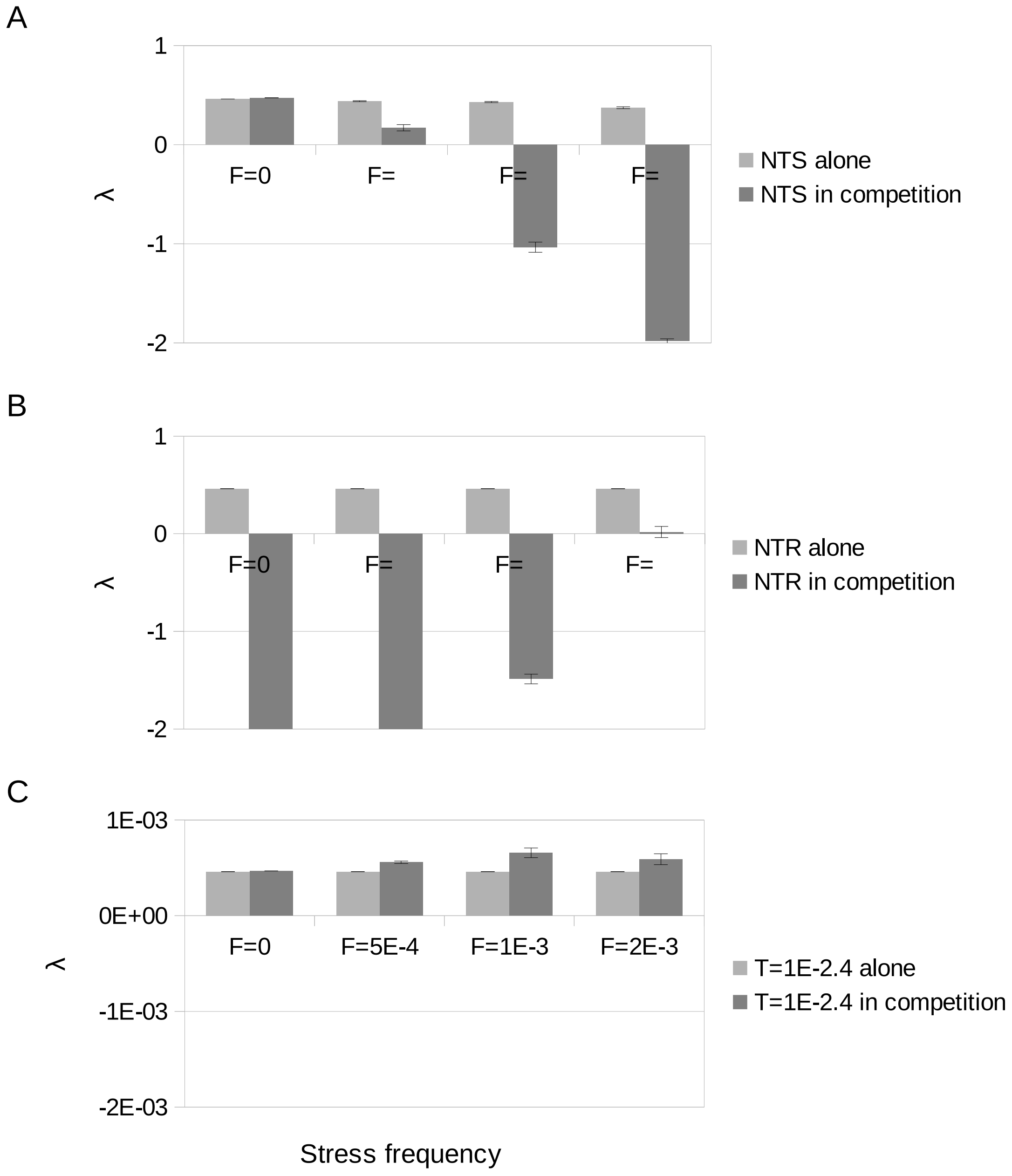


*Sup. Figure 2:* Stochastic growth rate (λ) of the NTR, NTS and the predominant genotype ($T_{max}$=$10^{-2.4}$$TU^{-1}$) alone and in competition. (A) NTS performs well alone despite being susceptible to the stress but is greatly counter selected when in competition in stochastic stressful environments. (B) NTR is not affected by the stresses and performs well alone. However, it is not competitive when stresses are infrequent. (C) The genotype with an intermediate transformation rate ($T_{max}$=$10^{-2.4}$$TU^{-1}$) performs well in all conditions.

λ

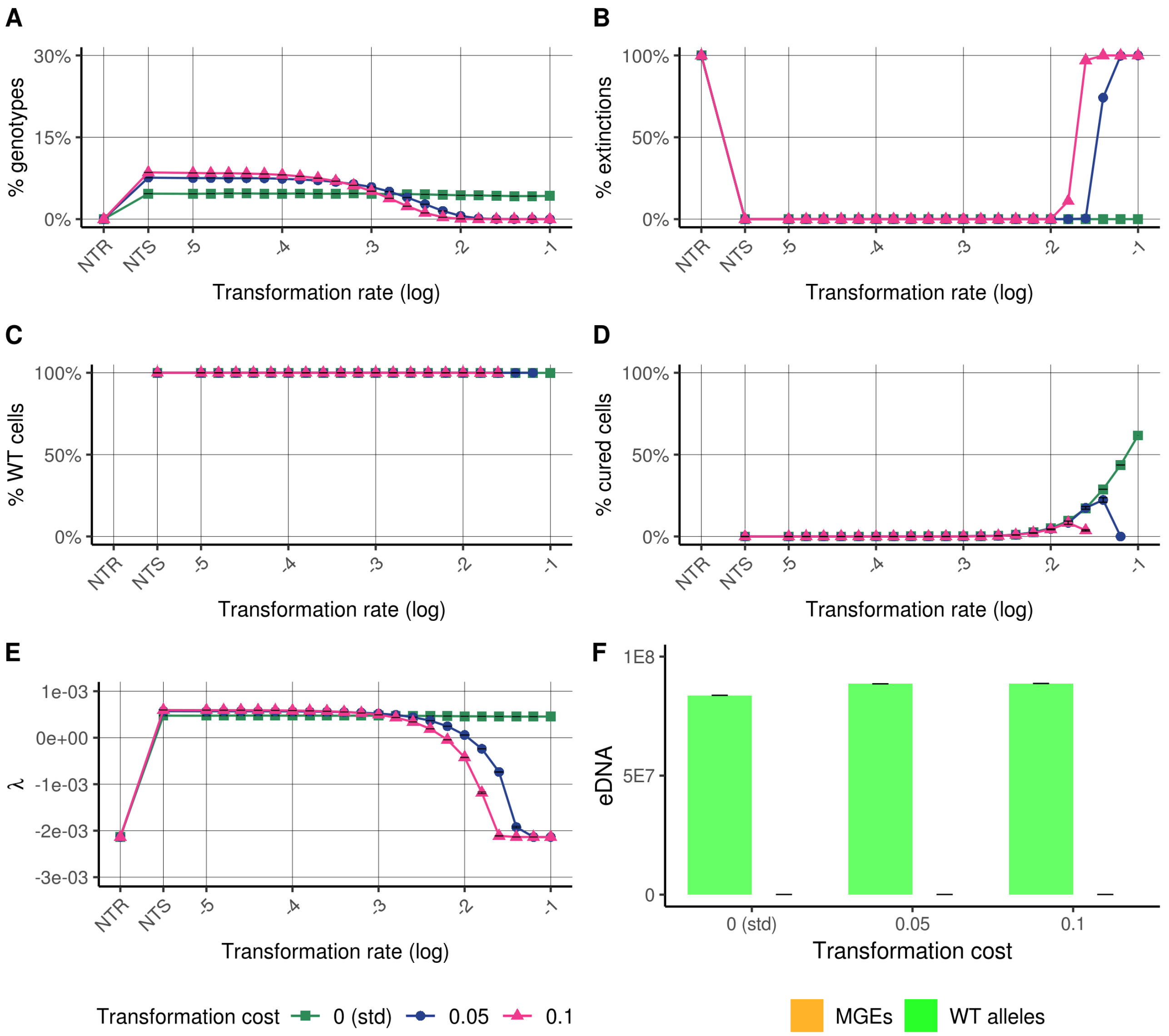


*Sup. Figure 3:* Sensitivity analysis to the transformation cost (probability of cell lysis during transformation events) in stress-free environment. Simulations with various transformation costs are compared to simulations with the standard parameters (std). (A) Distribution of the genotype at the end of the simulations. (B) Percentage of extinctions of the genotypes. (C) Percentage of wild type cells. (D) Percentage of wild type cells from cure. (E) Stochastic growth rate of each genotype (see methods). (F) Composition of the extracellular compartment, i.e. the number of eDNA molecules of each type, WT alleles and MGEs. Results are the mean and standard error of 500 simulations.

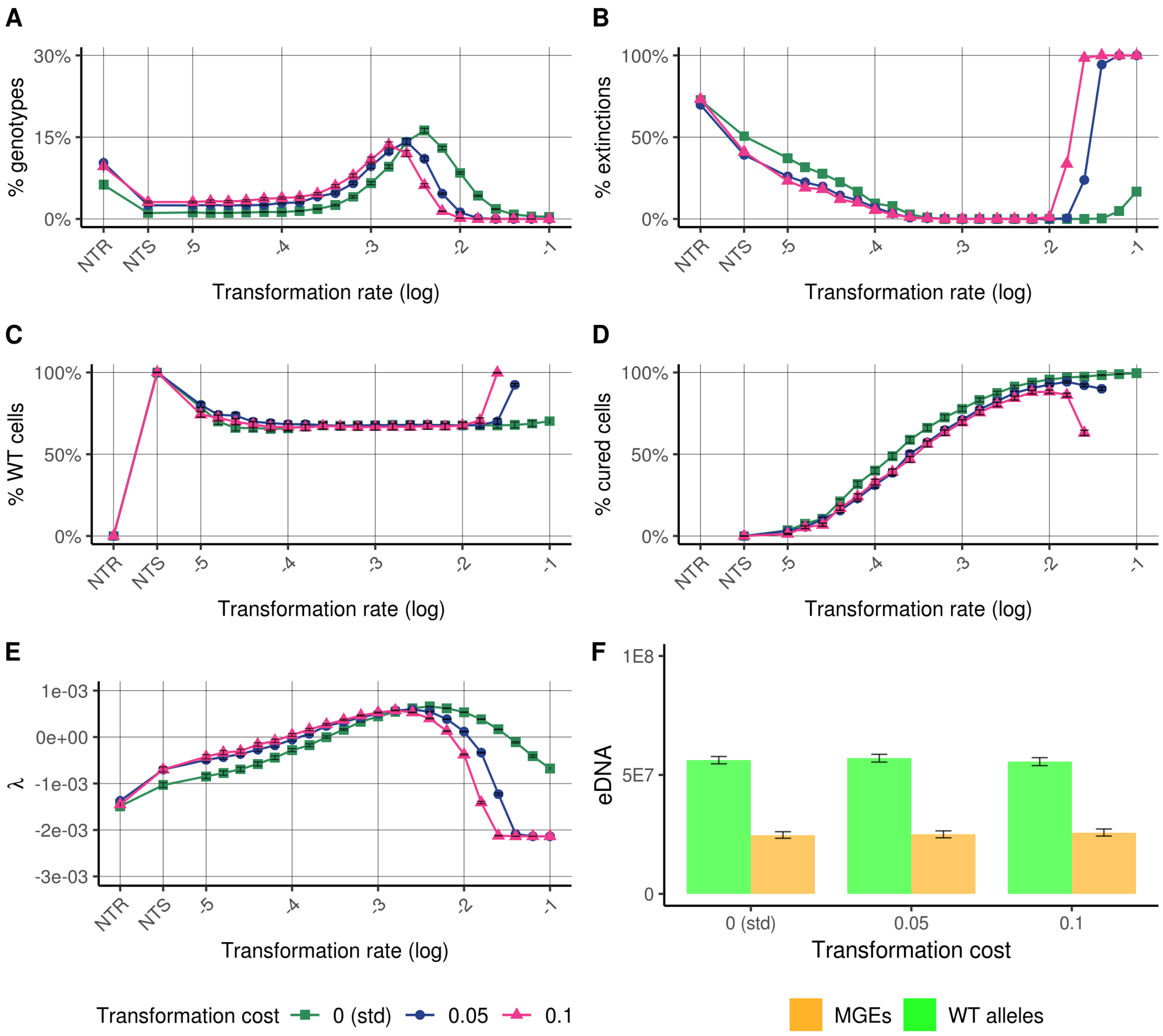


*Sup. Figure 4:* Sensitivity analysis to the transformation cost (probability of cell lysis during transformation events) in stressful environment (frequency of stress $F=10^{-3}UT^{-1}$). See Sup.figure 3 for more details about panels A to F.

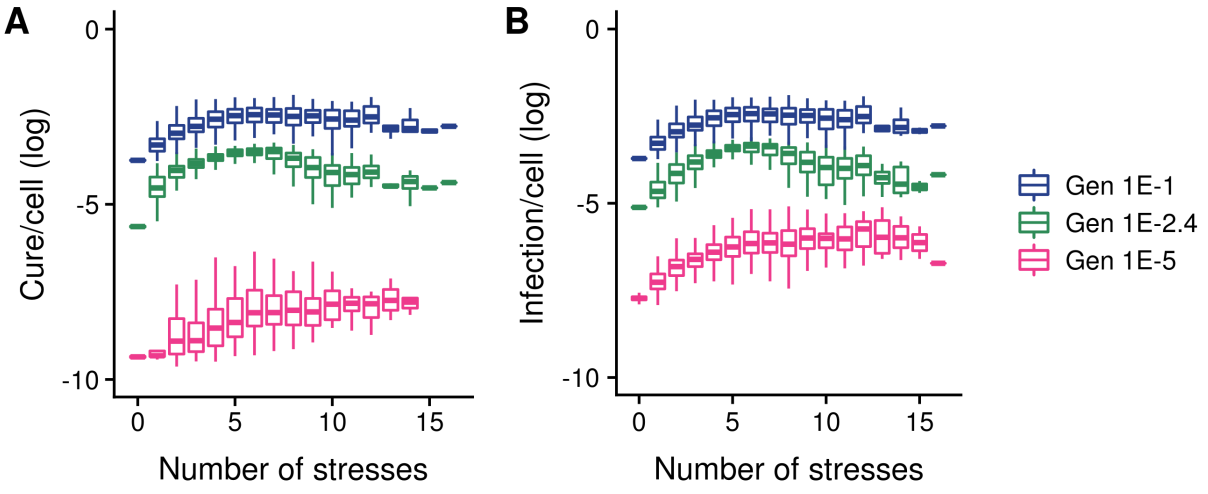


*Sup. Figure 5:* Types of transformation events per cell. The graphs represent frequency of cure (MGE infected ->WT) or infections (WT→MGE infected) per cell. Three genotypes with three transformation rates are plotted: a genotype with a very low transformation rate ($T_{max}$=$10^{-5}$), a genotype with an optimal transformation rate in a fluctuating environment ($T_{max}$=$10^{-2.4}$) and a genotype with a very high transformation rate ($T_{max}$=$10^{-1}$). Results presented are simulations with the standard parameters and the 4 environments presented in the main text.

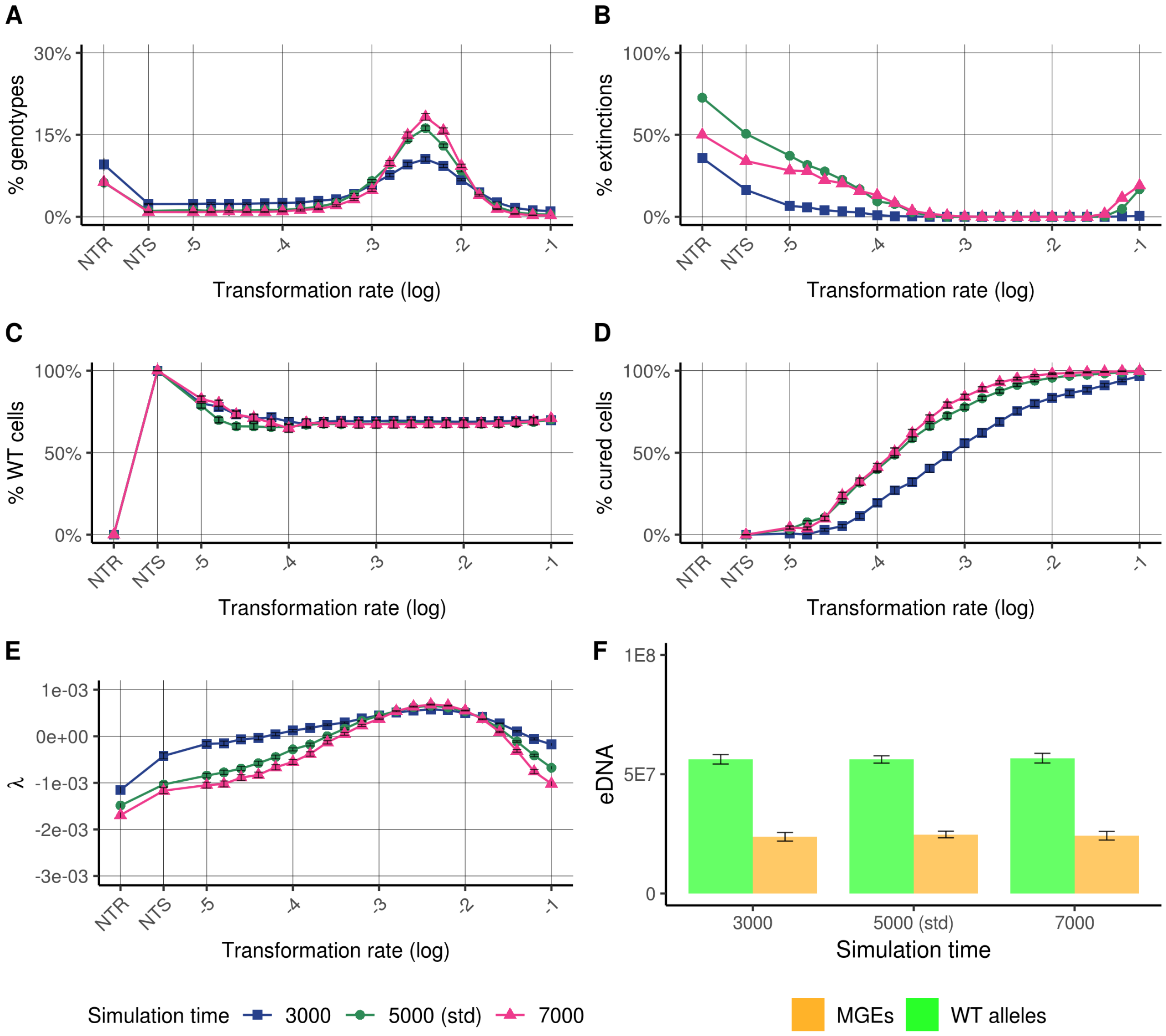


*Sup. Figure 6:* Sensitivity analysis to the duration of the simulations. See Sup.figure 3 for more details about panels A to F.

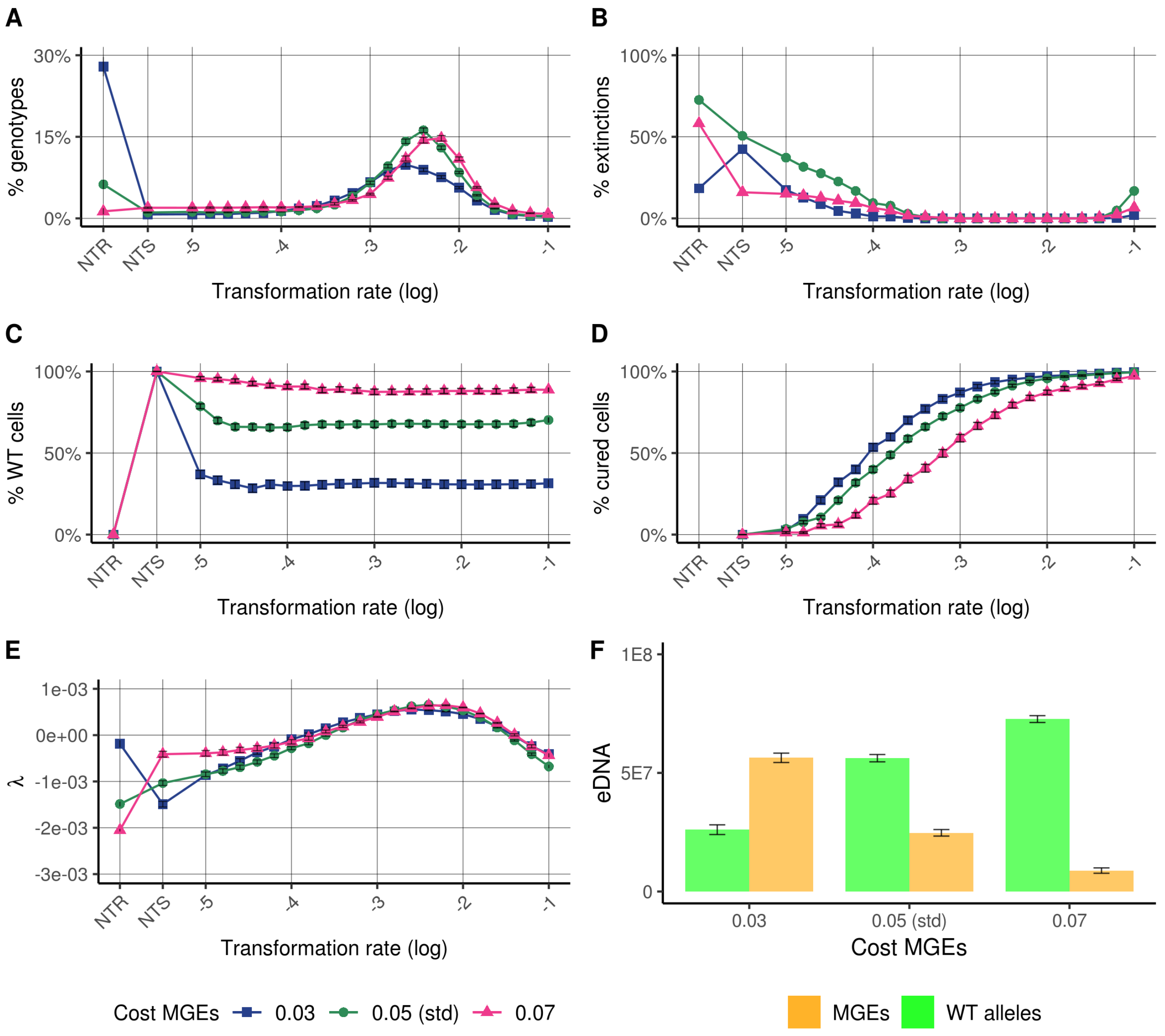


*Sup. Figure 7:* Sensitivity analysis to the cost of the resistance carried by Mobile Genetic Elements (cost  on the replication of cells $c_{MGE}$). See Sup.figure 3 for more details about panels A to F.

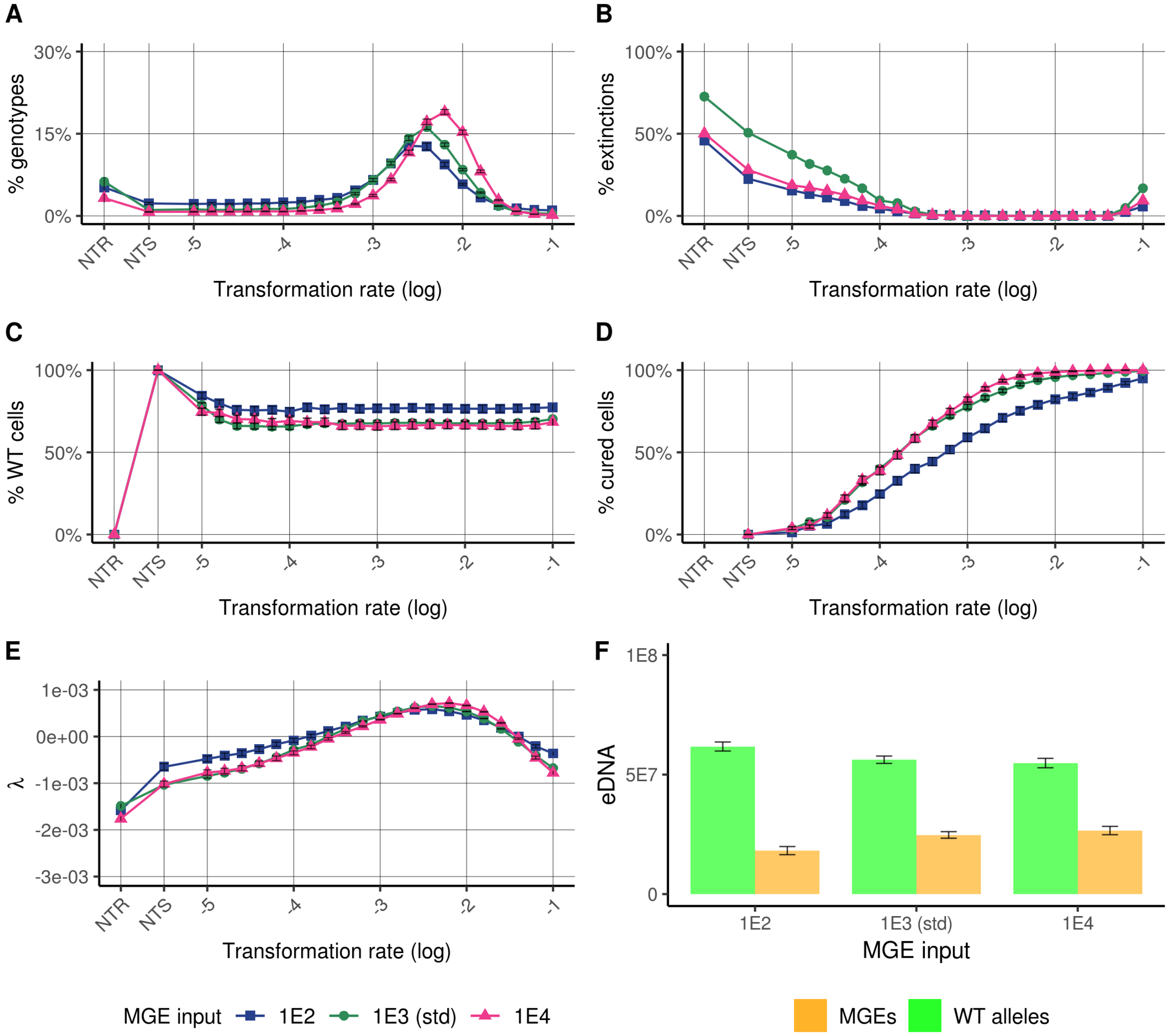


*Sup. Figure 8:* Sensitivity analysis to the input of Mobile Genetic Elements, i.e. the rate to which MGEs are added added to the extracellular environment, simulating a residual arrival from neighboring populations. See Sup.figure 3 for more details about panels A to F.

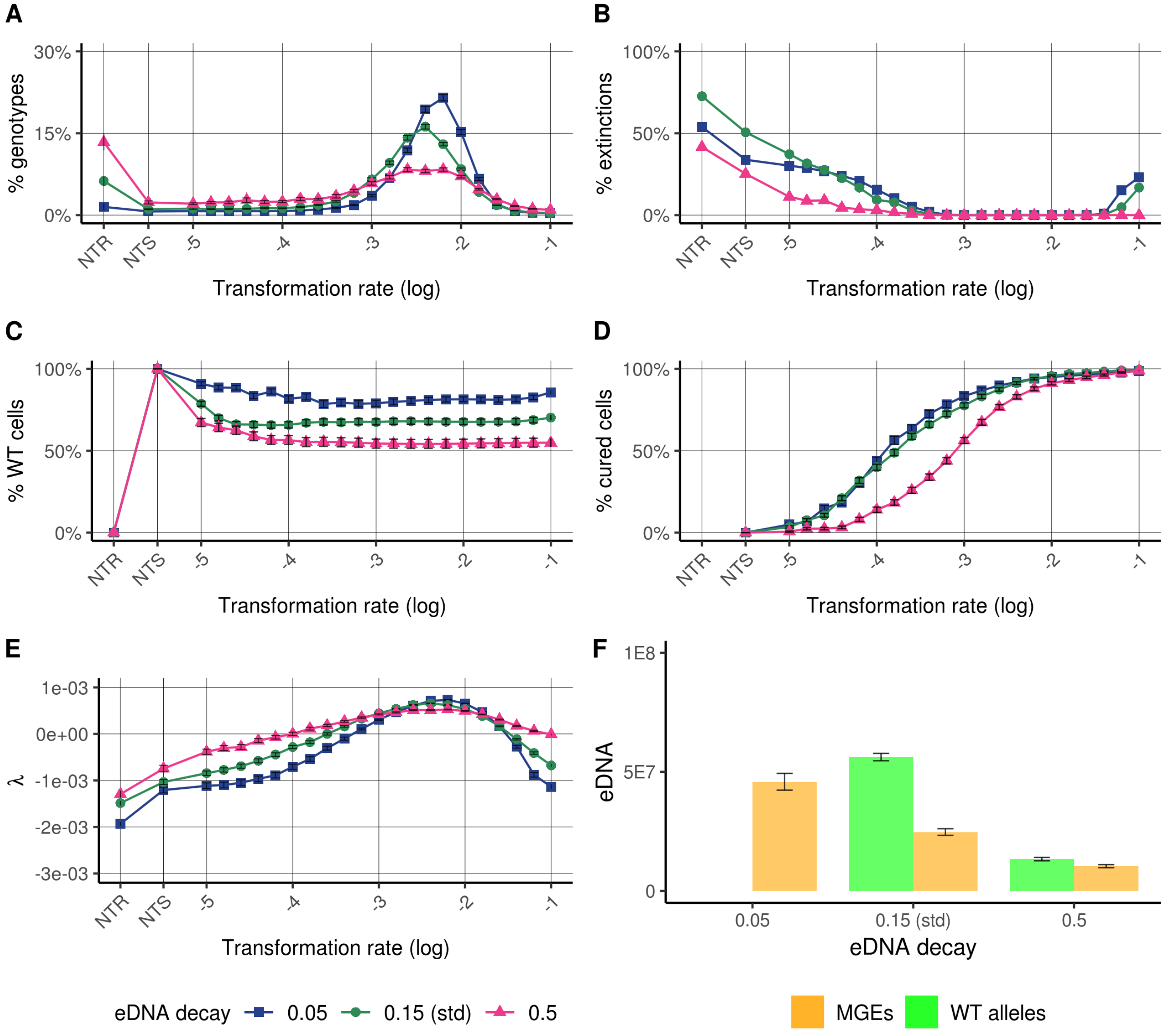


*Sup. Figure 9:* Sensitivity analysis to the eDNA decay. MGEs and WT alleles are degraded at the same rate. See Sup.figure 3 for more details about panels A to F.

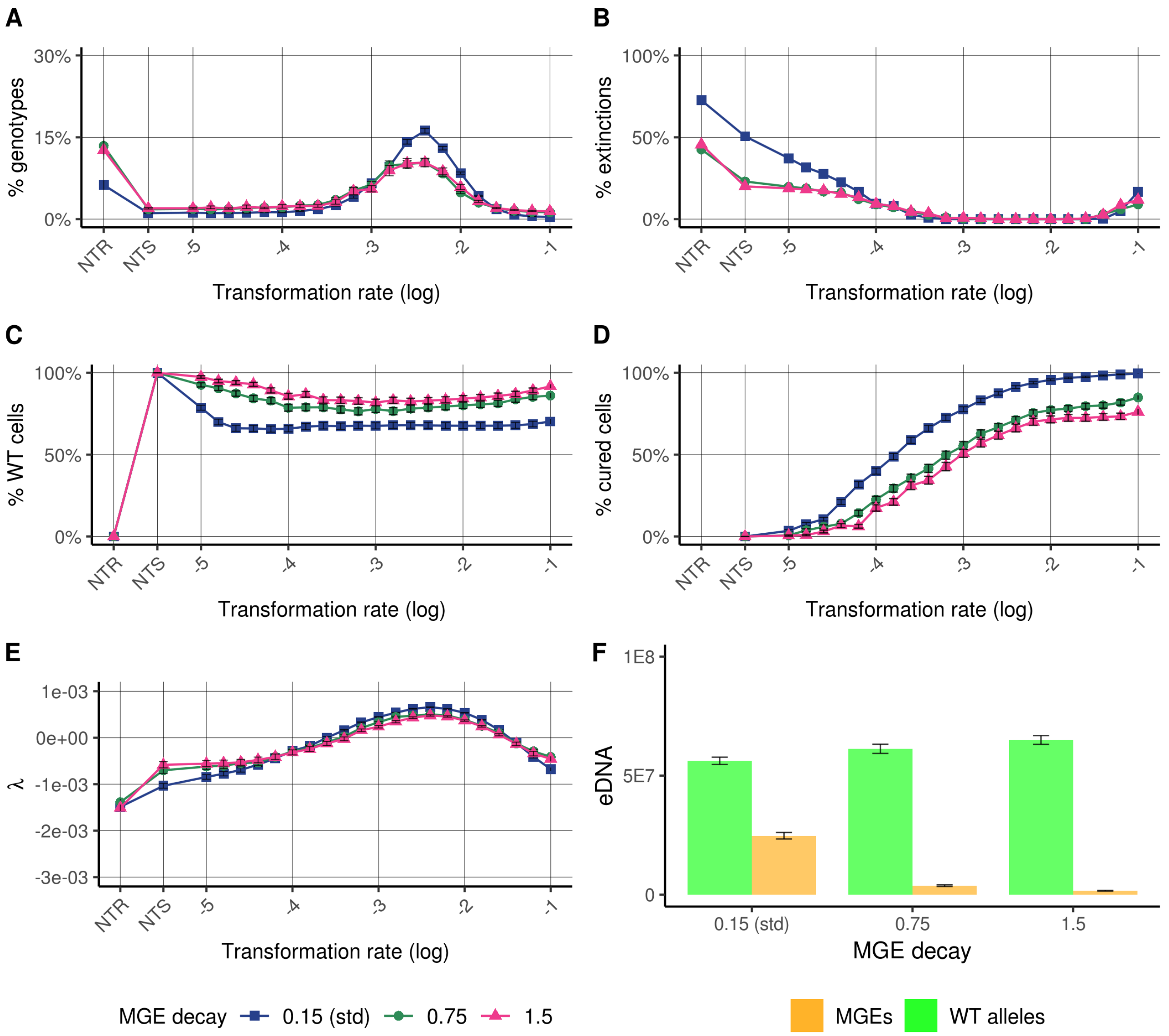


*Sup. Figure 10:* Sensitivity analysis to the extracellular MGE decay. MGEs are degraded at higher rates than WT alleles. Extracellular WT allele decay remains at 0.15. See Sup.figure 3 for more details about panels A to F.

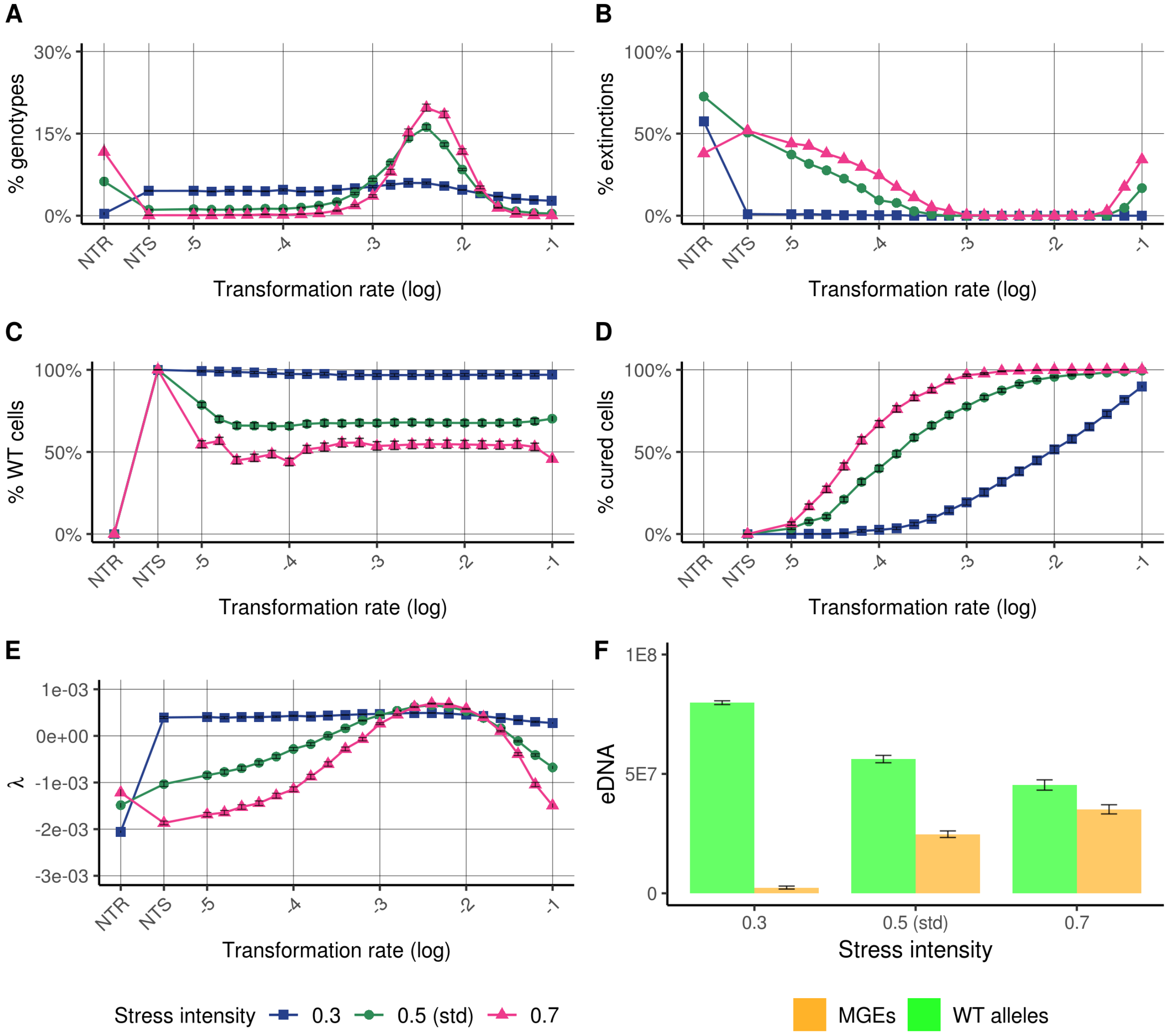


*Sup. Figure 11:* Sensitivity analysis to the mean stress intensity. See Sup.figure 3 for more details about panels A to F.

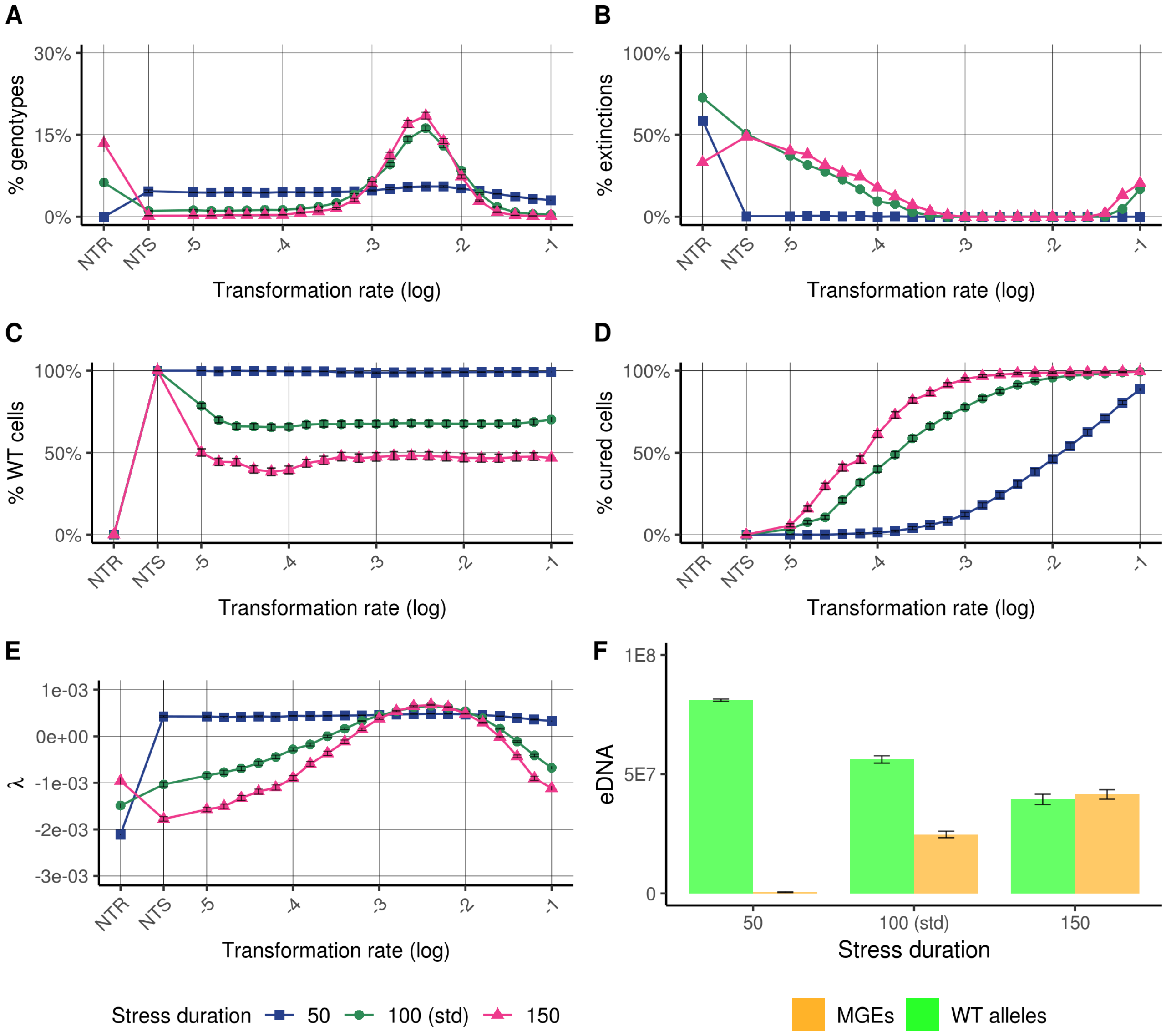


*Sup. Figure 12:* Sensitivity analysis to the m*ean stress duration. See Sup.figure 3 for* more details about panels A to F.
